# Methylation pattern of polymorphically imprinted *nc886* is not conserved across mammalia

**DOI:** 10.1101/2021.12.03.471165

**Authors:** Daria Kostiniuk, Hely Tamminen, Pashupati P. Mishra, Saara Marttila, Emma Raitoharju

**Affiliations:** Molecular Epidemiology, Faculty of Medicine and Health Technology, Tampere University, Tampere, Finland; Department of Clinical Chemistry, Faculty of Medicine and Health Technology, Tampere University, Tampere, Finland; Finnish Cardiovascular Research Centre, Faculty of Medicine and Health Technology, Tampere University, Tampere, Finland; Department of Clinical Chemistry, Fimlab Laboratories, Tampere, Finland; Gerontology Research Center, Tampere University, Finland; Tampere University Hospital, Tampere, Finland

**Keywords:** nc886, vtRNA2-1, primates, imprinting, DNA methylation

## Abstract

**Background:** In humans, the *nc886* locus is a polymorphically imprinted metastable epiallele. Periconceptional conditions have an effect on the methylation status of *nc886*, and further, this methylation status is associated with health outcomes in later life, in line with the Developmental Origins of Health and Disease (DOHaD) hypothesis. Animal models would offer opportunities to study the associations between periconceptional conditions, *nc886* methylation status and metabolic phenotypes further. Thus, we set out to investigate the methylation pattern of the *nc886* locus in non-human mammals.

**Data:** We obtained DNA methylation data from the data repository GEO for mammals, whose *nc886* gene included all three major parts of *nc886* and had sequency similarity of over 80% with the human *nc886*. Our final sample set consisted of DNA methylation data from humans, chimpanzees, bonobos, gorillas, orangutangs, baboons, macaques, vervets, marmosets and guinea pigs.

**Results:** In human data sets the methylation pattern of *nc886* locus followed the expected bimodal distribution, indicative of polymorphic imprinting. In great apes, we identified a unimodal DNA methylation pattern with 50% methylation level in all individuals and in all subspecies. In Old World monkeys, the between individual variation was greater and methylation on average was close to 60%. In guinea pigs the region around the *nc886* homologue was non-methylated. Results obtained from the sequence comparison of the CTCF binding sites flanking the *nc886* gene support the results on the DNA methylation data.

**Conclusions:** Our results indicate that unlike in humans, *nc886* is not a polymorphically imprinted metastable epiallele in non-human primates or in guinea pigs, thus implying that animal models are not applicable for *nc886* research. The obtained data suggests that the *nc886* region may be classically imprinted in great apes, and potentially also in Old World monkeys, but not in guinea pigs.

## Background

In mammalian genomic imprinting, only one parental allele is expressed, while gene expression from the other allele is suppressed in a parent-of-origin-dependent manner. Imprinting has been hypothesized to have evolved in response to a parent-offspring conflict, where the maternal and paternal genome differ in their interests regarding the supply of resources [1]. Paternal genomes favor the opportunistic strategy to enhance the growth of the developing offspring through the expression of growth-enhancing genes, while maternal genomes aim to conserve maternal resources over multiple pregnancies [2]. On the other hand, according to the trophoblast defense theory, suppression of genes that promote placental development or to activate genes that limit this process in oocytes are thought to protect the females from parthenogenetically activated oocytes and consequential malignant trophoblast [3,4]. The expression of imprinted genes in general has been associated with fetal and placental growth and suggested to a have a role in the development of cardiometabolic diseases in adulthood [2,4,5]. Imprinting arose relatively recently at most loci— while only a few imprinted genes in Eutherians are also imprinted in marsupials, to date no imprinting has been reported in the egg-laying monotreme mammals [6]. This supports the conflict-of-interest theory, as only in non-egg laying mammals the growing fetus directly consumes the maternal resources [4]. Genetic imprinting is best described in mice, and while the mouse is an informative proxy for human imprinted gene regulation, less than half of the 100 human imprinted genes have been shown to be similarly imprinted in mice (https://www.geneimprint.com/site/home). Distinct differences in placental evolution, physiology, and reproductive biology of the primate and murine groups may be responsible for these differences.

During gametogenesis and fertilization, the original DNA methylation pattern of imprinted genes is erased, and parent of origin-based methylation pattern is established. While most imprinted genes are located in clusters that are regulated by insulators or long noncoding RNAs [3], some unclustered imprinted genes can be regulated by differential promoter methylation [4]. Parental imprints are maintained after fertilization through these mechanisms despite extensive reprogramming of the mammalian genome [4]. A common feature of imprinted genes is insulators, such as CCCTC binding factor (CTCF) binding sites, which block the enhancers from interacting with gene promoters and/or act as barrier to the spread of transcriptionally repressive condensed chromatin [7].

The locus harbouring non coding 886 (*nc886*, also known as VTRNA2-1) in chromosome 5q31.1 is a unique example of imprinting, as it is the only known polymorphically imprinted locus across tissues in adult human population, where the polymorphism is not due to genetic variation [8–12]. However, in the placenta polymorphic imprinting is more common [13,14]. The *nc886* differentially methylated region (DMR) is 1.9-kb long, and its boundaries are marked by two CTCF binding sites [10,15]. This DMR has been shown to present maternal imprinting in ~75% of individuals in several populations [10,12,16]. This means that while in all individuals the paternal allele is unmethylated, in approximately 75% of individuals the maternal allele is methylated (individuals present a 50% methylation level at *nc886* locus) and in the remaining 25% of individuals the maternal allele is unmethylated (individuals present a 0% methylation level at the *nc886* locus).

The *nc886* gene codes for a 102nt long, non-coding RNA, which is then cleaved into two short RNAs (hsa-miR-886-3p/nc886-3p [23 nt] and hsa-miR-886-5p/nc886-5p [24–25 nt]) [17–19]. There is no consensus on whether the effects of *nc886* expression is mediated by the 102 nt long hairpin structure or the nc886-3p and −5p molecules, as the short molecules have been indicated to function as miRNAs, while the hairpin loop has been shown to inhibit protein kinase R (PKR) [17,20]. Expression of nc886 RNAs is strongly associated with the methylation status of the *nc886* locus. Individuals with non-methylated *nc886* present approximately two-fold levels of nc886 RNAs in blood, as compared to individuals with monoallelic methylation [16], thus implying allele-specific expression. However, allele-specific expression has not been experimentally shown, as the 102 nt transcript does not harbour SNPs [10].

The periconceptional environment has been suggested to affect DNA methylation patterns in maternal alleles[21], including the *nc886* epiallele [10,15,16]. Season of conception, maternal age and socioeconomic status have been linked to changes in the proportion of offspring with unmethylated maternal allele [10,15,16]. On the other hand, lower levels of *nc886* methylation have been linked to cleft palate [22], and a non-methylated *nc886* epiallele has been associated with an elevated childhood BMI [23]. The methylation status of this epiallele has also been associated with allergies [24], asthma [25], infections [26], and inflammation [27]. We and others have also shown that both the *nc886* methylation status and RNA expression are associated with indicators of glucose metabolism [16,28].

These results indicate that *nc886* could mediate the association between periconceptional conditions and later metabolic health, in line with the Developmental Origins of Health and Disease (DOHaD) hypothesis (aka the Barker hypothesis) [29]. More detailed analysis on periconceptional conditions and *nc886* methylation status and investigations between *nc886* and metabolic phenotypes, with less cofounding factors, would be needed to confirm this hypothesis. As carcinogenesis [9] and pluripotency induction [16] affect the DNA methylation pattern in *nc886* locus, *in vitro* work has its limitations. Animal models could be a feasible option for this research. Unfortunately, rodents do not harbor the *nc886* gene, limiting the use of traditional model organisms [10]. Thus, this study was set up to investigate 1) which animals have *nc886* gene, 2) whether this gene is surrounded by similar CTFC elements as the human homolog and 3) whether the methylation status of the *nc886* region suggest polymorphic imprinting in non-human mammals.

## Materials and methods

The presence of *nc886* gene was investigated in ensemble, in 65 amniota vertebrates Mercator-Pecan collection and 24 primates EPO-extended collection [30]. To select species for further investigation, we required the *nc886* gene have 80% sequence similarity with the human homolog and to present the sequences for nc886-3p and nc886-5p RNAs, as well as the loop structure, previously shown to mediate the binding of PKR [17,31] (S1 and S2 Figs). The existence and sequence similarity of the centromeric (chr5:135415115-135415544) and telomeric CTCF (chr5:135418124-135418523) binding site flanking *nc886* gene were also investigated in species shown to harbor intact *nc886* gene. If homologous CTCF-binding sites were not discovered, CTCFBSDB 2.0 [32] was utilized to predict possible non-homologous sites. Interactions of the *nc886* flanking CTCF-sites were also investigated using K562 CTCF ChIA-PET Interactions data and Hi-C data in genome browser [33] and Hi-C data in 3D Genome Browser from HUVEC and K562 cell lines [34].

For species harboring the *nc886* gene, we investigated the Gene Expression Omnibus (GEO) repository [35] for available DNA methylation data, with both general and binomial name of the species. DNA methylation data was available in apes from chimpanzees (*Pan troglodytes*, n=83; GSE136296 [36] and n=5; GSE41782 [37]), bonobos (*Pan paniscus*, n=6; GSE41782 [37]), gorillas (*Troglodytes gorilla*, n=6; GSE41782 [37]) and orangutangs (*Pongo spp*., n=6; GSE41782[37]). In Old World monkeys, data was obtained from baboons (*Papio spp.* n=28; GSE103287 [38]), rhesus macaques (*Macaca mulatta,* n=10; GSE103287 [38]), vervets (*Chlorocebus aethiops*, n=10; GSE103287 [38]) and in New World monkeys, from marmosets (*Callithrix jacchus*, n=6; GSE103287 [38]). From primates, only data from blood or femur was utilized. In addition to primates, we obtained DNA methylation data from guinea pig hippocampus (*Cavia porcellus,* n=36; GSE109765 [39]). As reference, we utilized data from human (*Homo sapiens*) blood (n=1658; GSE105018 [40]), femur (n=48; GSE64490 [41]) and hippocampus (n=33; GSE72778 [42]).

Methylation profiling data obtained by high throughput sequencing (guinea pigs, GSE109765) was processed as follows. Quality of the paired-end reads in all the samples was assessed using FastQC [43] and MultiQC [44]. Paired-end fastq files were trimmed using Trimmomatic-0.39 with a sliding window of size 4 set to remove bases with phred score lower than 20 [45]. The trimmed samples were analyzed using Bismark-0.23.0 tools [46]. The reads were aligned to the guinea pig genome (cavPor3). Duplicate alignments, which can arise for example by PCR amplification, were removed. Methylation information was extracted from the alignment result files using Bismark’s methylation extractor. DNA methylation values for CpGs inside the gene were first inspected and then a wider region (±2000nt) around the gene was investigated.

Primate DNA methylation data from GSE41782, GSE105018, GSE64490, GSE72778 (profiled with Illumina 450K) and GSE136296 (profiled with Illumina EPIC) were available as processed data and was used as such. Primate DNA methylation data from GSE103271, GSE103280, GSE103286, which are subseries of GSE103332 (profiled with Illumina EPIC), were available as raw data, and were normalized by using minfi quantile normalization for each species separately.

From primate data, the 14 CpGs in the *nc886* DMR previously reported to show bimodal methylation pattern in humans were retrieved [10,16]. In all the primate species, for which methylation data was available, the sequence on the binding site of the Illumina probes was investigated. Only data from sites with the CG-sequence intact in the species in question were further utilized. (S1 File). Similar process was repeated for 50 probes in paternally expressed 10 (*PEG10*) previously shown to be imprinted [47]. *PEG10* was used as evolutionally conserved reference for a classically imprinted gene [6].

## Results and Discussion

### *nc886* gene in non-human mammals

Human *nc886* has been suggested to be an evolutionally young gene, producing a 102 nt long RNA, which is then ineffectively cleaved to two miRNA-like RNAs [17]. In line with previous reports [10]*, nc886* gene, with intact short RNA coding sequences and the hairpin loop, can be found in primates, in guinea pig (*Cavia porcellus*), Eurasian red squirrel (*Sciurus vulgaris*), and Alpine marmot (*Marmota marmota*), with two of the latter having insertions in the centromeric end of the gene (Fig 1, Fig 2, S1 Fig, S1 Table). In humans, and all the other species where *nc886* was found, *nc886* is located between *TGFB1* and *SMAD5*.

**Fig 1.**
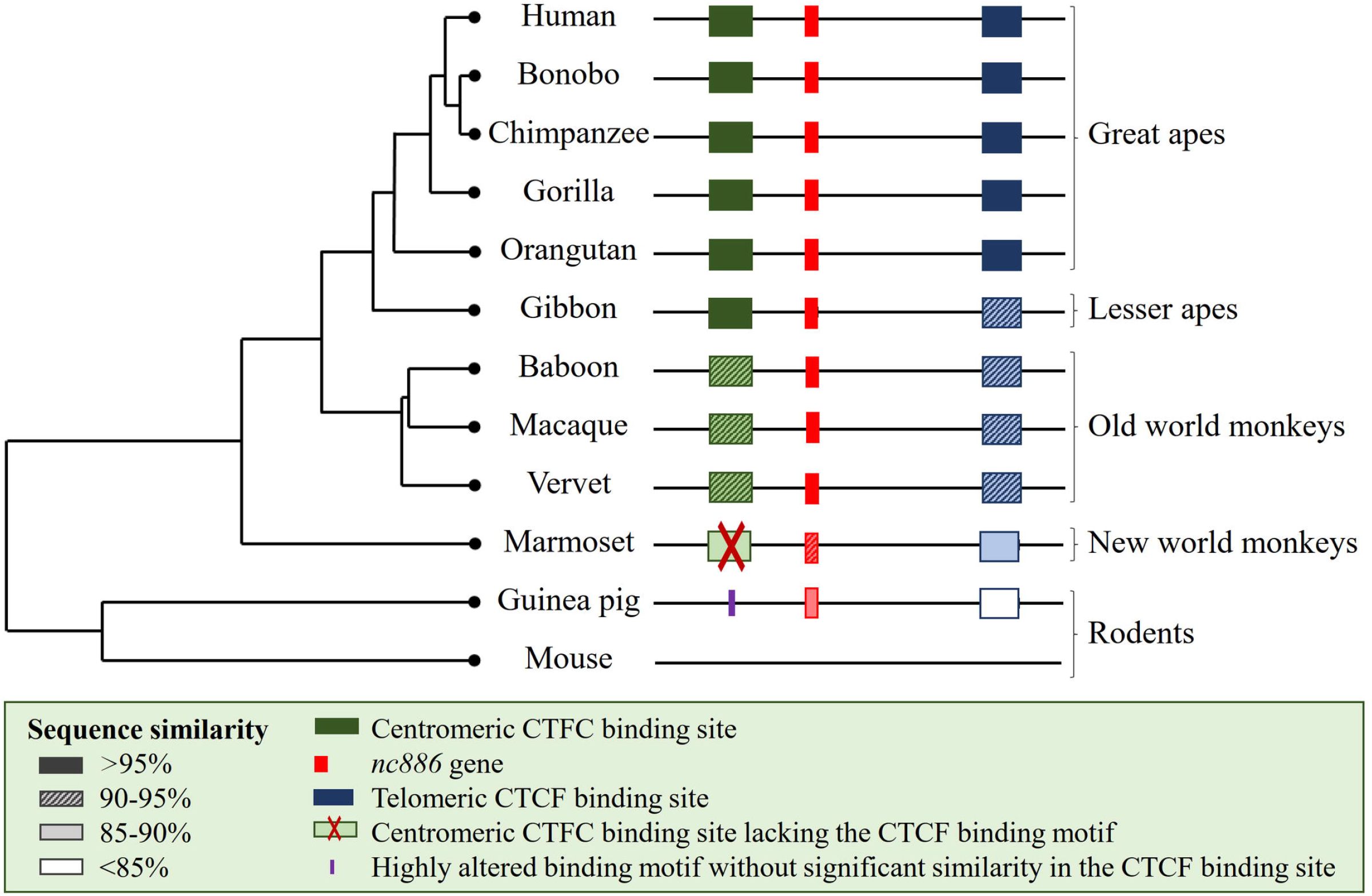
Schematic presentation of the similarity of the *nc886* gene and flanking CTCF binding sites. Guinea pigs lack the whole centromeric flanking CTCF binding site, whilst the marmosets have a region with sequence similarity, but lack the binding sequence of the CTCF. Mouse genome does not contain either the *nc886* gene or the CTCF binding sites flanking it. For detailed sequence comparisons, see S1 and S2 Figs.

**Fig 2.**
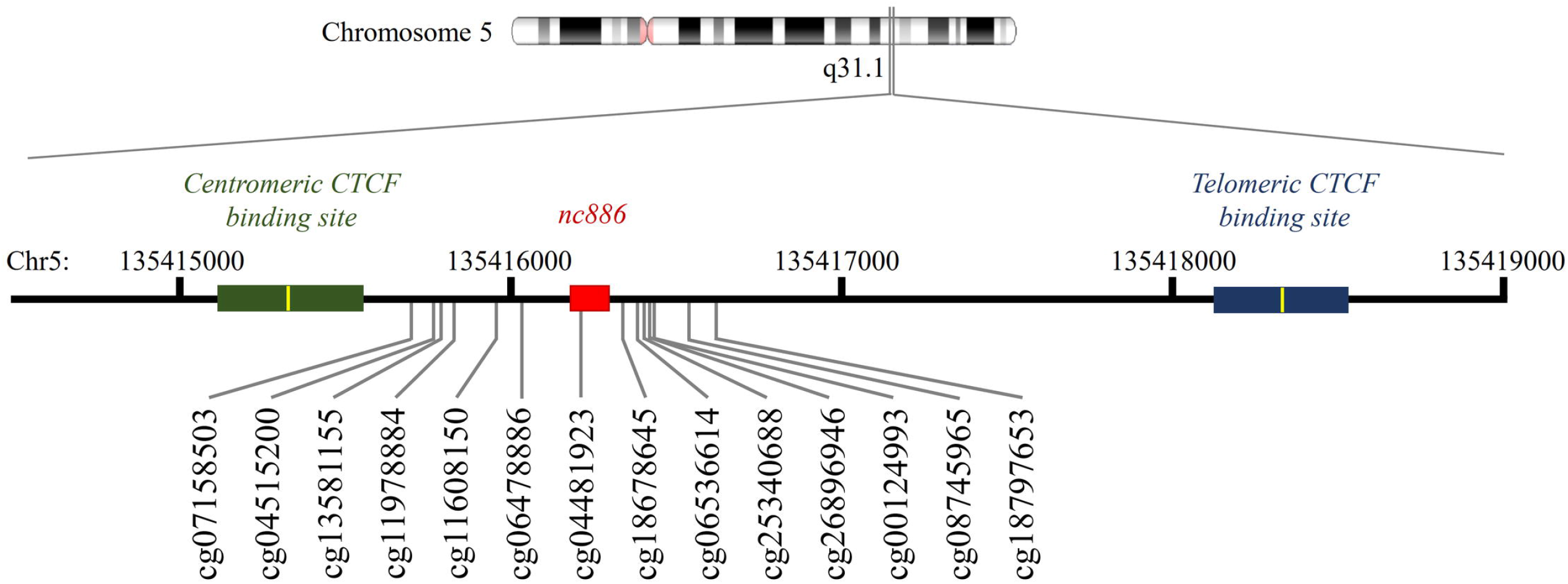
Schematic presentation of the *nc886* locus. In red, the *nc886* gene (chr5:135416180-135416287), in green the centromeric CTCF binding site (chr5:135415115-135415544) and in blue the telomeric CTFC binding site (chr5:135418124-135418523). The binding motif of the CTCF sites are presented in yellow and the 14 CpGs with bimodal methylation pattern in human cohorts are presented according to their genomic location (GRCh37/hg19).

Upon further inspection of primate *nc886* gene, almost 100% sequence similarity was identified in apes (*hominoidea*). The sequence similarity was high (over 97%) between humans and the investigated Old World monkeys (*Cercopithecidae*), while less (over 90%) similarity can be seen between humans and New World monkeys (*Ceboidea*) and even less (84-89%) between humans and tarsiers (*Tarsiidae*) or lemurs (*Lemuroidea*). Sequences coding for the short nc886 RNAs are identical within Old World anthropoids (*Catarrhini).* Differences between human and New World monkey and prosimian *nc886* sequences can be found both in the short nc886 RNA and the hairpin coding regions, but the most significant differences can be found in the centromeric end of the gene (Fig 1 and S2 Fig).

### CTCF binding site sequence in mammals with *nc886* gene

In humans, the *nc886* locus is flanked by two CTCF binding sites (Fig 2) [10]. The telomeric CTCF binding site can be found in most vertebrates, with the binding motif being identical in all species showed to have the *nc886* gene (S1 Table). The telomeric CTCF binding site was shown to interact with a CTCF binding site (chr5:135222814-135223707) locating near the *IL9* gene in the CTCF ChIA-PET data, the interaction is also supported by the Hi-C data from cell lines (S3 Fig). This prediction is in line with our previous finding indicating that genetic polymorphisms near *IL9* gene are associated with the expression of nc886 RNAs [16]. This CTCF binding site locating near *IL9* can be found in all primates (S1 Table). Together these results suggest that the evolutionally conserved telomeric CTCF binding site of *nc886* interacts with CTCF binding sites near *IL9* gene, possibly forming a topologically associating domain (TAD), or an interaction within one (sub-TAD), and bringing the suggested enhancer area near the *nc886* gene [48]. On the contrary, the centromeric CTCF binding site is present only in primates and even in primates, the binding sequence cannot be identified in marmosets. As CTCF binding sites can act as barriers to the spread of transcriptionally repressive condensed chromatin [7], presence, or absence, of the CTCF binding site may be associated with the methylation status of *nc886* locus in different species. For the centromeric CTCF binding site no interactions were detected according to CTCF ChIA-PET data. There are also changes in the CTFC binding motif in gorillas (position 9), in all Old World monkeys (position 4) and also in New World monkeys (position 14) (S1 Table). According to the CTCFBSDB 2.0, the guinea pig genome does not harbor any non-homologous predicted CTCF binding sites in the centromeric side of the *nc886* gene.

### Guinea pigs present a non-methylated *nc886* locus

One of the aims of this study was to investigate whether the information gathered from the *nc886* gene and DMR from cell culture and population studies could be supplemented with research on animal models. In line with a previous report [10] we identified this gene only in primates, guinea pigs and few members of the squirrel family, of which guinea pig was the most promising candidate as a model organism. In data from Constantinof et al. [39], in guinea pig hippocampi (n=36) the whole *nc886/vtRNA2-1* gene was non-methylated (S4 Fig). The surrounding *nc886* region (Scaffold DS562872.1: 24,622,179-24,622,280) +/− 2000 nt was mostly unmethylated, with only 2% of the reads in the region being methylated. It should be noted that the number of reads in the region in the data utilized was low (max number of reads=17, average number of reads=7). *nc886* methylation pattern in human hippocampi presented the expected bimodal distribution, and thus the discovered methylation pattern in the guinea pig hippocampi was most likely not due to the selection of tissue (S4 Fig). This identified lack of methylation in the *nc886* locus is compatible with the absence of the telomeric CTCF binding site, as CTCF binding sites can delineate the boundaries of an imprinted region [49]. These results thus suggest that guinea pigs are not suitable model organisms for the investigation of establishment of *nc886* methylation status.

### Imprinted *nc886* region in great apes

The blood of chimpanzees, gorillas, bonobos and orangutans presented beta-values close to 0.5 with a unimodal distribution in the *nc886* region (Fig 3, S2 Table). This methylation pattern closely resembles the methylation pattern in the known maternally imprinted gene *PEG10* (S5 Fig). The methylation levels are also very similar to those presented in humans with monoallelic methylation (Fig 3). In these data sets, that include more than 110 apes, we did not identify any individuals with methylation level close to 0 in the *nc886* locus, whereas in humans 25% of the population present a methylation level close to 0 at this locus [10,16]. If the prevalence on non-methylated individuals in apes was similar to humans, already 11 individuals would present at least one non-methylated individual with 95% probability. Of individual species, we had the largest dataset for chimpanzees (n=83 in GSE136296 and n=5 in GSE41782). Again, assuming the same proportion of non-methylated individuals as in humans (25%), probability of not identifying any non-methylated chimpanzees in a population of 88 individuals is extremely low, 1.01*10^−11^. These results imply that the *nc886* locus is not polymorphically imprinted in apes.

**Figure 3.**
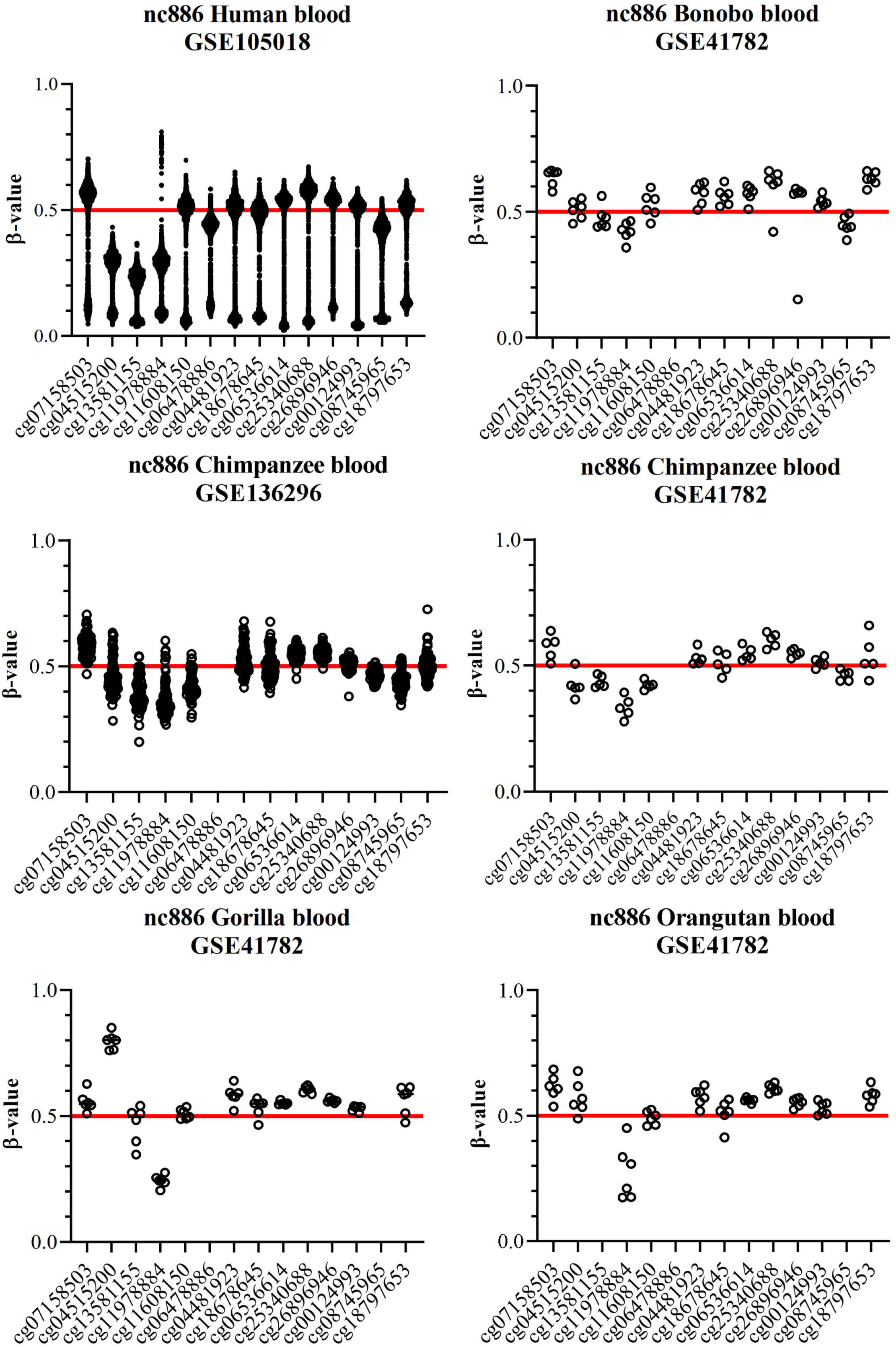
Blood DNA methylation beta values in *nc886* region in great apes (Hominidae). In each graph, one dot represents one individual. In humans a bimodal methylation pattern can be detected, in line with the expected population distribution of individuals with monoallelic methylation (75% of the population, methylation level ~0.5) and non-methylated individuals (25% of the population, methylation level close to 0) [10,16]. All of the other species present a unimodal methylation pattern, with all individuals having methylation beta-values near 0.5 across the *nc886* locus. Number of individuals is 1658 for humans (GSE105018), 6 for bonobos (GSE41782), 83 and 5 for chimpanzees (GSE136296 and GSE41782, respectively), 6 for gorillas (GSE41782) and 6 for orangutans (GSE41782). For schematic representation of the CpG sites in the *nc886* locus, see Fig 2.

Within species, median of standard deviation of the probe methylation of *nc886* is around 0.04, which is comparable to that seen in *PEG10* in apes (median of SD 0.03) and in human blood (0.02), indicating good data quality (S2 Table). In all apes the methylation levels of *nc886* region resembled those seen in *PEG10,* which is an evolutionally conserved maternally imprinted gene [6]. It is thus reasonable to suggest that *nc886* could be classically imprinted in other great apes, excluding humans.

### *nc886* region methylation patterns in Old World monkeys

The patterns of *nc886* region methylation are very similar in all of the Old World monkeys. The median methylation level is close to 0.60, higher as compared to apes. The interindividual variation is larger, especially in baboons, than in apes, with the median SD within a probe being 0.11. The SD within a probe is also higher in probes locating in *PEG10* in baboons, where the median of probe SD is 0.05 (S2 Table). As the between individual variation in methylation levels of a known evolutionally conserved imprinted gene is also higher, this suggests a technical bias in the data, potentially due to the use of Illumina Infinium 450K and EPIC methylation assays, that are designed for humans. All methylation data available for Old World monkeys was from femur, but as the methylation pattern of *nc886* has been reported to be constant across tissues in humans [12] and in the human reference data set form femur samples both *nc886* and *PEG10* present similar methylation patterns as in blood (Fig 4 and S6 Fig), this phenomenon most likely is not caused by the tissue of origin. Regardless of the precise methylation levels of the Old World monkeys, in the 48 individuals we did not identify any presenting a non-methylated methylation pattern in *nc886* region, probability of which is 1.0*10^−6^, when assuming similar distribution as in humans. This implies that similar to non-human great apes, the *nc886* locus is not polymorphically imprinted in Old World monkeys.

**Figure 4.**
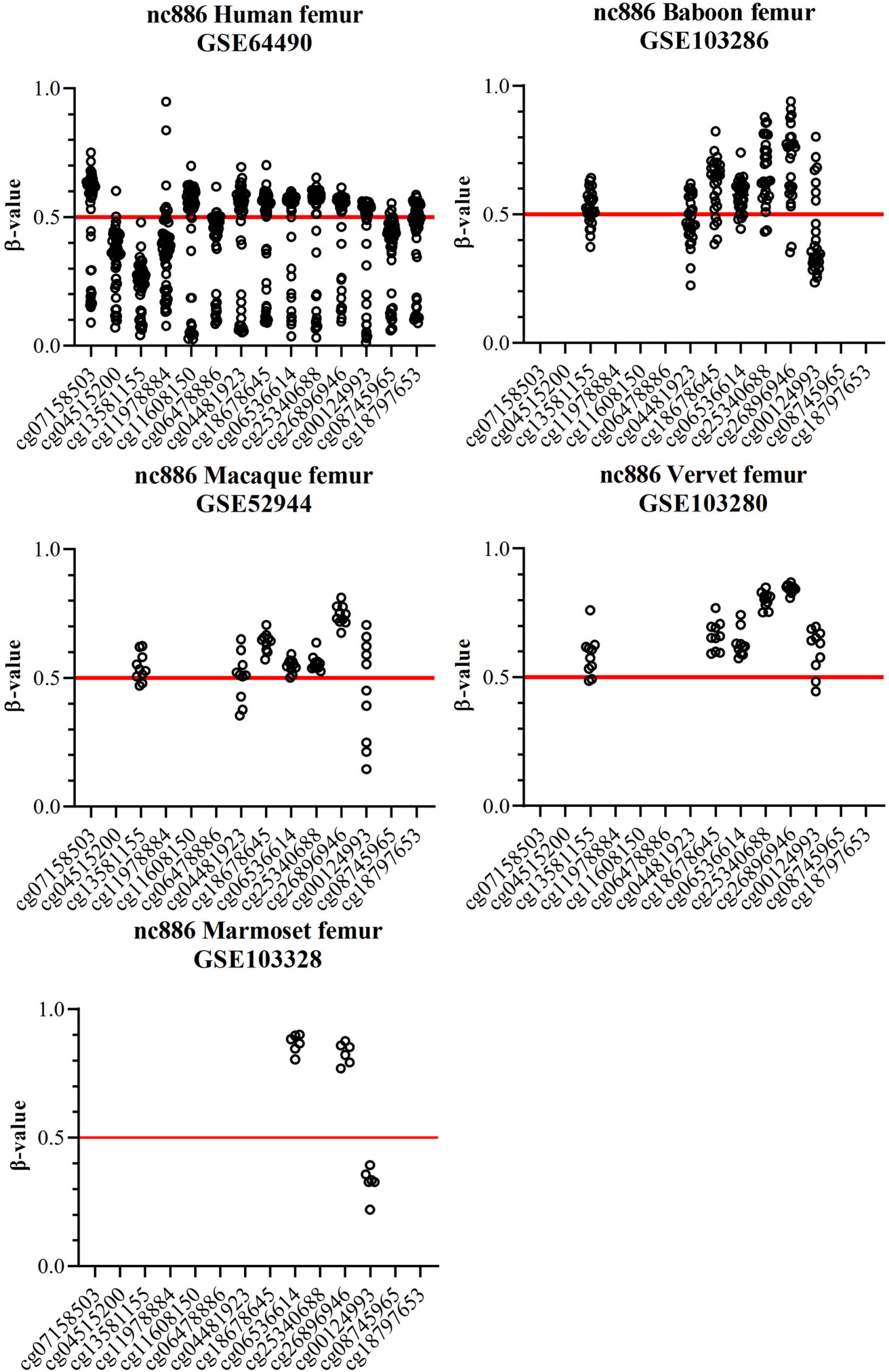
Femur DNA methylation beta values in the *nc886* region in humans and in monkeys. In humans a bimodal methylation pattern can be detected, similar to blood and in line with the expected population distribution of individuals with monoallelic methylation (75% of the population, methylation level ~0.5) and non-methylated individuals (25% of the population, methylation level close to 0) [10,16]. In Old World monkeys the between individual variation is bigger than in apes. No individuals presenting a non-methylated *nc886* region are detected. In marmosets only 3 probes were considered to provide reliable methylation values and none of them present methylation levels near 0.5. Number of individuals is 48 for humans (GSE64490), 28 for baboons (GSE103287), 10 for macaques (GSE103287), 10 for vervets (GSE103287) and 6 for marmosets (GSE103287). For schematic representation of the CpG sites in the *nc886* locus, see Fig 2.

### nc886 region methylation patterns in New World monkeys

Data from the six marmosets is not conclusive, as we found that only 3 of the probes in *nc886* region bound to areas with no great sequence differences. The methylation levels of two of these probes were around 0.8 and one 0.32, showing no indications of imprinting, while the median of methylation beta values in the probes locating in the *PEG10* is 0.46 in marmosets (Fig 4 and S6 Fig, S2 Table). The lack of imprinting of any kind in marmosets is further supported by the finding that they lack the centromeric CTCF binding motif, which is thought to have an important role in insulating the DMR [10]. To draw conclusions on the methylation status of *nc886* in New World monkeys in general, methylation data from more species would be needed.

### Limitations of the study

Our study is purely descriptive in nature. In guinea pigs, the shallowness of the sequencing data limits the ability to make conclusions of the methylation pattern. In non-human primates, utilizing methylation arrays that have been designed for humans raises questions on data quality, especially in marmosets. Concerns over data quality are however mitigated by the observed methylation pattern in well-established imprinted gene, *PEG10*, as well as the consistency of observed results across different species and data sets. In addition, while a methylation level of 0.50 implies allele specific methylation, we were not able to confirm this due to lack of suitable data.

## Conclusions

We describe here an analysis on the methylation status of *nc886* region in non-human mammals. A genetic locus, with more than 80% similarity to human *nc886* gene can be found in primates, guinea pigs and some members of the squirrel family. We obtained DNA methylation data for 8 different non-human primate species and for guinea pigs, and in none of these species could we observe methylation pattern indicative of similar polymorphic imprinting, as could be observed, and has been reported [10,12,16], in humans. The observed methylation pattern in apes and in Old World monkeys implies that the *nc886* region might be classically imprinted, although these findings have to be interpreted with caution, as apart from chimpanzees the sample number were low, and in Old World monkeys the variation between individuals was notable. In guinea pigs, the most feasible potential model organism of those harboring the *nc886* locus, the data indicated that the locus is completely unmethylated. It is noteworthy that only primates, whose genome also contained the centromeric CTCF binding sequence flanking the *nc886* gene, had methylation levels indicative of genetic imprinting.

In conclusion, we were unable to identify an animal model suited to study the establishment of the methylation pattern of the polymorphically imprinted metastable epiallele *nc886*. Further studies on how this kind of unusual metastable developed, and how it links to the periconceptional conditions and later life health traits, are thus restricted to *in vitro* and population studies.

## Declarations

**Ethics approval**

Datasets used in this study were retrieved from Gene Expression Omnibus (GEO, www.ncbi.nlm.nih.gov/geo/) repository. For human data sets, informed consent or “Consent for Autopsy” were given by all participants. For animal studies all protocols were approved by the local ethical committees and/or the samples were collected during standard veterinary checks or routine necropsies. Details can be found from the original publications [36–42].

## Consent for publication

Not applicable

## Availability of data and materials

All methylation data utilized are available in GEO, under accession numbers GSE136296, GSE41782, GSE103287, GSE109765, GSE105018, GSE64490 and GSE72778.

## Author Contributions

S.M. and E.R. designed the research; D.K., H.T., P.P.M., S.M. and E.R. performed research; S.M. and E.R. provided resources; D.K., H.T., P.P.M., S.M. and E.R. curated data; S.M. and E.R wrote the original draft manuscript; D.K., H.T., P.P.M., S.M. and E.R. reviewed and edited the manuscript. Funding was acquired by S.M and E.R. All authors have read and agreed upon the published version of the manuscript.

## Acknowledgements

The authors wish to thank Jooseppi Järvinen (MSc) for his contribution to the probability calculations, Nina Mononen (PhD) and Binisha Mishra (MSc) for their valuable input on the manuscript and to acknowledge all the researchers who contributed towards the collection of datasets utilized in this study.

## Supporting information

**S1 Fig. Sequence alignment of the *nc886* gene in the 65 amniota vertebrates Mercator-Pecan.** To state that a species presents the *nc886* we required 80% sequence similarity with the human *nc886* and presence of both nc886-5p and nc886-3p short RNAs and the hairpin loop present in the 102nt long nc886 RNA. Species above the dashed line were considered to harbour the full *nc886* gene. Note: Sequence alignment figures are presented in the direction of the gene. Species with no identified alignment in this region were excluded from the figure (Sauropsids, opossum (*Monodelphis domestica*) and platypus (*Ornithorhynchus anatinus*)).

**S2 Fig. Sequence alignment of the *nc886* gene in the 24 primates in EPO-extended collection.** Sequence similarity to human *nc886* gene decreases as evolutionary distance increases and greatest diverge is seen in the centromeric end of the gene. Note: Sequence alignment figures are presented in the direction of the gene.

**S3 Fig. Predicted interaction between *nc886* telomeric CTCF binding site and CTCF binding site located near IL9.** A) ChIA-PET data and in HI-C data in Genome browser cell lines and in 3D genome browser. B) HUVEC and C) K562 cell lines. The suggested sub-TAD has been indicated with a black square and interactions in HI-C data have been circumscribed. The telomeric CTCF binding site is located at chr5:135418124-135418523 and the CTCF binding site near *IL9* at chr5:135223050-135223420.

**S4 Fig. Methylation of *nc886* region in hippocampi of A) guinea pigs (GSE109765) and B) humans (GSE64509).** In (A) each circle presents a CpG site with measured methylation level and each line one individual guinea pig (n=36). The number of reads per site is low (on average 7) but none of the samples provide data that would indicate anything but non-methylated DNA methylation status around the guinea pig *nc886* gene. In human hippocampi (B) the methylation pattern of *nc886* presents the expected binomial methylation pattern, which is in line with reported frequency of 25% non-methylated individuals and 75% individuals with monoallelic methylation in a population.

**S5 Fig. Methylation pattern of Paternally expressed 10 (*PEG10*) in blood of great apes.** Only probes locating in sites with no clear sequence differences as compared to the human *PEG10* sequence are shown.

**S6 Fig. Methylation pattern of Paternally expressed 10 (*PEG10*) in femur bone of humans and Old and New World monkeys.** Only probes locating in sites with no clear sequence differences as compared to the human *PEG10* sequence are shown.

**S1 File. Sequence alignment of the *nc886* region between human, bonobo, chimpanzee, gorilla, orangutang, gibbon, baboon, macaque, vervet and marmoset.** The sequence comparison covers the region containing the 14 CpG sites presenting bimodal methylation pattern in humans (chr5:135415593-135416666). These CpG sites are highlighted in yellow.

**S1 Table. Sequence similarities of *nc886* gene and its centromeric and telomeric CTCF binding sites, and the CTCF binding site located near Il9 gene in primates, marmot, squirrel, guinea pig and in mouse investigated from NCBI BLAST and ensemble**.

**S2 Table. Methylation medians and standard deviations of the PEG10 and *nc886* regions and individual probes in the *nc886* region in primates.** Medians and standard deviations in human *nc886* are not comparable to other species, as it is known to demonstrate bimodal methylation pattern in humans.

## References

1. Moore T, Haig D. Genomic imprinting in mammalian development: a parental tug-of-war. Trends in Genetics. 1991. doi:10.1016/0168-9525(91)90230-N

2. Smith FM, Garfield AS, Ward A. Regulation of growth and metabolism by imprinted genes. Cytogenetic and Genome Research. 2006. doi:10.1159/000090843

3. Varmuza S, Mann M. Genomic imprinting - defusing the ovarian time bomb. Trends in Genetics. 1994;10. doi:10.1016/0168-9525(94)90212-7

4. Barlow DP, Bartolomei MS. Genomic imprinting in mammals. Cold Spring Harbor Perspectives in Biology. 2014;6. doi:10.1101/cshperspect.a018382

5. Millership SJ, Van de Pette M, Withers DJ. Genomic imprinting and its effects on postnatal growth and adult metabolism. Cellular and Molecular Life Sciences. 2019. doi:10.1007/s00018-019-03197-z

6. Renfree MB, Suzuki S, Kaneko-Ishino T. The origin and evolution of genomic imprinting and viviparity in mammals. Philosophical Transactions of the Royal Society B: Biological Sciences. 2013. doi:10.1098/rstb.2012.0151

7. Lewis A, Reik W. How imprinting centres work. Cytogenetic and Genome Research. 2006. doi:10.1159/000090818

8. Treppendahl MB, Qiu X, Søgaard A, Yang X, Nandrup-Bus C, Hother C, et al. Allelic methylation levels of the noncoding VTRNA2-1 located on chromosome 5q31.1 predict outcome in AML. Blood. 2012. doi:10.1182/blood-2011-06-362541

9. Romanelli V, Nakabayashi K, Vizoso M, Moran S, Iglesias-Platas I, Sugahara N, et al. Variable maternal methylation overlapping the nc886/vtRNA2-1 locus is locked between hypermethylated repeats and is frequently altered in cancer. Epigenetics. 2014. doi:10.4161/epi.28323

10. Carpenter BL, Zhou W, Madaj Z, DeWitt AK, Ross JP, Grønbæk K, et al. Mother–child transmission of epigenetic information by tunable polymorphic imprinting. Proceedings of the National Academy of Sciences of the United States of America. 2018. doi:10.1073/pnas.1815005115

11. Zink F, Magnusdottir DN, Magnusson OT, Walker NJ, Morris TJ, Sigurdsson A, et al. Insights into imprinting from parent-of-origin phased methylomes and transcriptomes. Nature Genetics. 2018. doi:10.1038/s41588-018-0232-7

12. Carpenter BL, Remba TK, Thomas SL, Madaj Z, Brink L, Tiedemann RL, et al. Oocyte age and preconceptual alcohol use are highly correlated with epigenetic imprinting of a noncoding RNA (nc886). Proceedings of the National Academy of Sciences of the United States of America. 2021;118. doi:10.1073/pnas.2026580118

13. Hanna CW, Peñaherrera MS, Saadeh H, Andrews S, McFadden DE, Kelsey G, et al. Pervasive polymorphic imprinted methylation in the human placenta. Genome Research. 2016;26. doi:10.1101/gr.196139.115

14. Sanchez-Delgado M, Court F, Vidal E, Medrano J, Monteagudo-Sánchez A, Martin-Trujillo A, et al. Human Oocyte-Derived Methylation Differences Persist in the Placenta Revealing Widespread Transient Imprinting. PLoS Genetics. 2016;12. doi:10.1371/journal.pgen.1006427

15. Silver MJ, Kessler NJ, Hennig BJ, Dominguez-Salas P, Laritsky E, Baker MS, et al. Independent genomewide screens identify the tumor suppressor VTRNA2-1 as a human epiallele responsive to periconceptional environment. Genome Biology. 2015. doi:10.1186/s13059-015-0660-y

16. Marttila S, Viiri LE, Mishra PP, Kühnel B, Matias-Garcia PR, Lyytikäinen L-P, et al. Methylation status of nc886 epiallele reflects periconceptional conditions and is associated with glucose metabolism through nc886 RNAs. Clinical Epigenetics. 2021;13. doi:10.1186/s13148-021-01132-3

17. Fort RS, Garat B, Sotelo-Silveira JR, Duhagon MA. vtRNA2-1/nc886 produces a small RNA that contributes to its tumor suppression action through the microRNA pathway in prostate cancer. Non-coding RNA. 2020. doi:10.3390/ncrna6010007

18. Kong L, Hao Q, Wang Y, Zhou P, Zou B, Zhang Y xiang. Regulation of p53 expression and apoptosis by vault RNA2-1-5p in cervical cancer cells. Oncotarget. 2015. doi:10.18632/oncotarget.4948

19. Lee K, Kunkeaw N, Jeon SH, Lee I, Johnson BH, Kang GY, et al. Precursor miR-886, a novel noncoding RNA repressed in cancer, associates with PKR and modulates its activity. RNA. 2011. doi:10.1261/rna.2701111

20. Lee YS. A Novel Type of Non-coding RNA, nc886, Implicated in Tumor Sensing and Suppression. Genomics & Informatics. 2015. doi:10.5808/gi.2015.13.2.26

21. Steegers-Theunissen RPM, Twigt J, Pestinger V, Sinclair KD. The periconceptional period, reproduction and long-term health of offspring: The importance of one-carbon metabolism. Human Reproduction Update. 2013. doi:10.1093/humupd/dmt041

22. Gonseth S, Shaw GM, Roy R, Segal MR, Asrani K, Rine J, et al. Epigenomic profiling of newborns with isolated orofacial clefts reveals widespread DNA methylation changes and implicates metastable epiallele regions in disease risk. Epigenetics. 2019. doi:10.1080/15592294.2019.1581591

23. van Dijk SJ, Peters TJ, Buckley M, Zhou J, Jones PA, Gibson RA, et al. DNA methylation in blood from neonatal screening cards and the association with BMI and insulin sensitivity in early childhood. International Journal of Obesity. 2018. doi:10.1038/ijo.2017.228

24. Yu S, Zhang R, Liu G, Yan Z, Hu H, Yu S, et al. Microarray analysis of differentially expressed microRNAs in allergic rhinitis. American Journal of Rhinology and Allergy. 2011. doi:10.2500/ajra.2011.25.3682

25. Suojalehto H, Lindström I, Majuri ML, Mitts C, Karjalainen J, Wolff H, et al. Altered microRNA expression of nasal mucosa in long-term asthma and allergic rhinitis. International Archives of Allergy and Immunology. 2014. doi:10.1159/000358486

26. Sharbati J, Lewin A, Kutz-Lohroff B, Kamal E, Einspanier R, Sharbati S. Integrated microrna-mrna-analysis of human monocyte derived macrophages upon mycobacterium avium subsp. hominissuis infection. PLoS ONE. 2011. doi:10.1371/journal.pone.0020258

27. Asaoka T, Sotolongo B, Island ER, Tryphonopoulos P, Selvaggi G, Moon J, et al. MicroRNA signature of intestinal acute cellular rejection in formalin-fixed paraffin-embedded mucosal biopsies. American Journal of Transplantation. 2012. doi:10.1111/j.1600-6143.2011.03807.x

28. Lin CH, Lee YS, Huang YY, Tsai CN. Methylation status of vault rna 2-1 promoter is a predictor of glycemic response to glucagon-like peptide-1 analog therapy in type 2 diabetes mellitus. BMJ Open Diabetes Research and Care. 2021;9. doi:10.1136/bmjdrc-2020-001416

29. Barker DJP, Osmond C. INFANT MORTALITY, CHILDHOOD NUTRITION, AND ISCHAEMIC HEART DISEASE IN ENGLAND AND WALES. The Lancet. 1986. doi:10.1016/S0140-6736(86)91340-1

30. Kent WJ, Sugnet CW, Furey TS, Roskin KM, Pringle TH, Zahler AM, et al. The Human Genome Browser at UCSC. Genome Research. 2002;12. doi:10.1101/gr.229102

31. Bou-Nader C, Gordon JM, Henderson FE, Zhang J. The search for a PKR code— differential regulation of protein kinase R activity by diverse RNA and protein regulators. RNA. 2019. doi:10.1261/rna.070169.118

32. Ziebarth JD, Bhattacharya A, Cui Y. CTCFBSDB 2.0: A database for CTCF-binding sites and genome organization. Nucleic Acids Research. 2013;41. doi:10.1093/nar/gks1165

33. Rosenbloom KR, Sloan CA, Malladi VS, Dreszer TR, Learned K, Kirkup VM, et al. ENCODE Data in the UCSC Genome Browser: Year 5 update. Nucleic Acids Research. 2013;41. doi:10.1093/nar/gks1172

34. Wang Y, Song F, Zhang B, Zhang L, Xu J, Kuang D, et al. The 3D Genome Browser: A web-based browser for visualizing 3D genome organization and long-range chromatin interactions. Genome Biology. 2018;19. doi:10.1186/s13059-018-1519-9

35. Barrett T, Wilhite SE, Ledoux P, Evangelista C, Kim IF, Tomashevsky M, et al. NCBI GEO: Archive for functional genomics data sets - Update. Nucleic Acids Research. 2013;41. doi:10.1093/nar/gks1193

36. Guevara EE, Lawler RR, Staes N, White CM, Sherwood CC, Ely JJ, et al. Age-associated epigenetic change in chimpanzees and humans. Philosophical Transactions of the Royal Society B: Biological Sciences. 2020;375. doi:10.1098/rstb.2019.0616

37. Hernando-Herraez I, Prado-Martinez J, Garg P, Fernandez-Callejo M, Heyn H, Hvilsom C, et al. Dynamics of DNA Methylation in Recent Human and Great Ape Evolution. PLoS Genetics. 2013;9. doi:10.1371/journal.pgen.1003763

38. Housman G, Quillen EE, Stone AC. Intraspecific and interspecific investigations of skeletal DNA methylation and femur morphology in primates. American Journal of Physical Anthropology. 2020;173. doi:10.1002/ajpa.24041

39. Constantinof A, Boureau L, Moisiadis VG, Kostaki A, Szyf M, Matthews SG. Prenatal Glucocorticoid Exposure Results in Changes in Gene Transcription and DNA Methylation in the Female Juvenile Guinea Pig Hippocampus Across Three Generations. Scientific Reports. 2019;9. doi:10.1038/s41598-019-54456-9

40. Hannon E, Knox O, Sugden K, Burrage J, Wong CCY, Belsky DW, et al. Characterizing genetic and environmental influences on variable DNA methylation using monozygotic and dizygotic twins. PLoS Genetics. 2018. doi:10.1371/journal.pgen.1007544

41. Horvath S, Mah V, Lu AT, Woo JS, Choi OW, Jasinska AJ, et al. The cerebellum ages slowly according to the epigenetic clock. Aging. 2015;7. doi:10.18632/aging.100742

42. Horvath S, Langfelder P, Kwak S, Aaronson J, Rosinski J, Vogt TF, et al. Huntington’s disease accelerates epigenetic aging of human brain and disrupts DNA methylation levels. Aging. 2016;8. doi:10.18632/aging.101005

43. Andrews S. FastQC - A quality control tool for high throughput sequence data. http://www.bioinformatics.babraham.ac.uk/projects/fastqc/. Babraham Bioinformatics. 2010.

44. Ewels P, Magnusson M, Lundin S, Käller M. MultiQC: Summarize analysis results for multiple tools and samples in a single report. Bioinformatics. 2016;32. doi:10.1093/bioinformatics/btw354

45. Bolger AM, Lohse M, Usadel B. Trimmomatic: A flexible trimmer for Illumina sequence data. Bioinformatics. 2014;30. doi:10.1093/bioinformatics/btu170

46. Krueger F, Andrews SR. Bismark: A flexible aligner and methylation caller for Bisulfite-Seq applications. Bioinformatics. 2011;27. doi:10.1093/bioinformatics/btr167

47. Hernandez Mora JR, Tayama C, Sánchez-Delgado M, Monteagudo-Sánchez A, Hata K, Ogata T, et al. Characterization of parent-of-origin methylation using the Illumina Infinium MethylationEPIC array platform. Epigenomics. 2018;10. doi:10.2217/epi-2017-0172

48. Kentepozidou E, Aitken SJ, Feig C, Stefflova K, Ibarra-Soria X, Odom DT, et al. Clustered CTCF binding is an evolutionary mechanism to maintain topologically associating domains. Genome Biology. 2020;21. doi:10.1186/s13059-019-1894-x

49. Dowen JM, Fan ZP, Hnisz D, Ren G, Abraham BJ, Zhang LN, et al. Control of cell identity genes occurs in insulated neighborhoods in mammalian chromosomes. Cell. 2014;159. doi:10.1016/j.cell.2014.09.030

