## Supplementary figures and images for "Methylation pattern of polymorphically imprinted *nc886* is not conserved across mammalia"

### S1 Figure

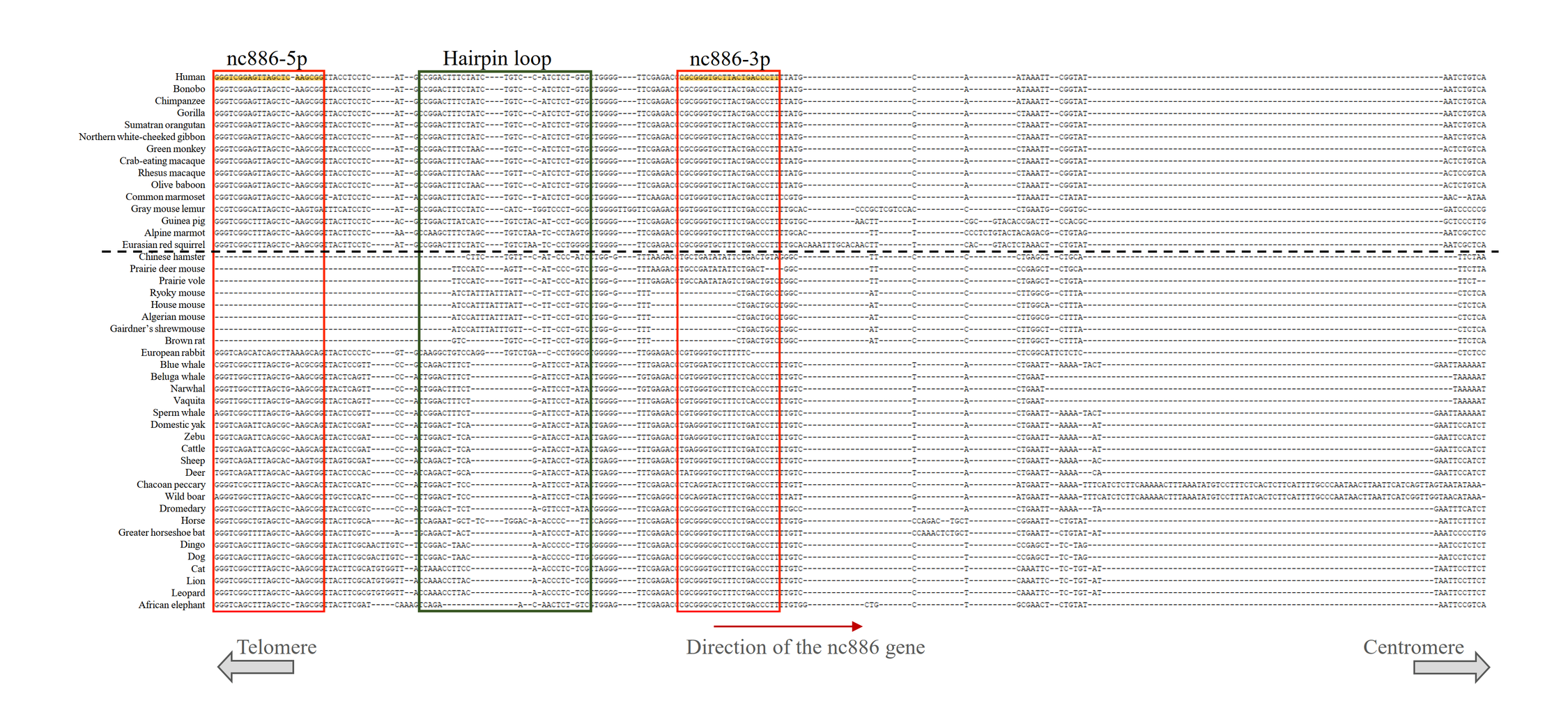

### S2 Figure

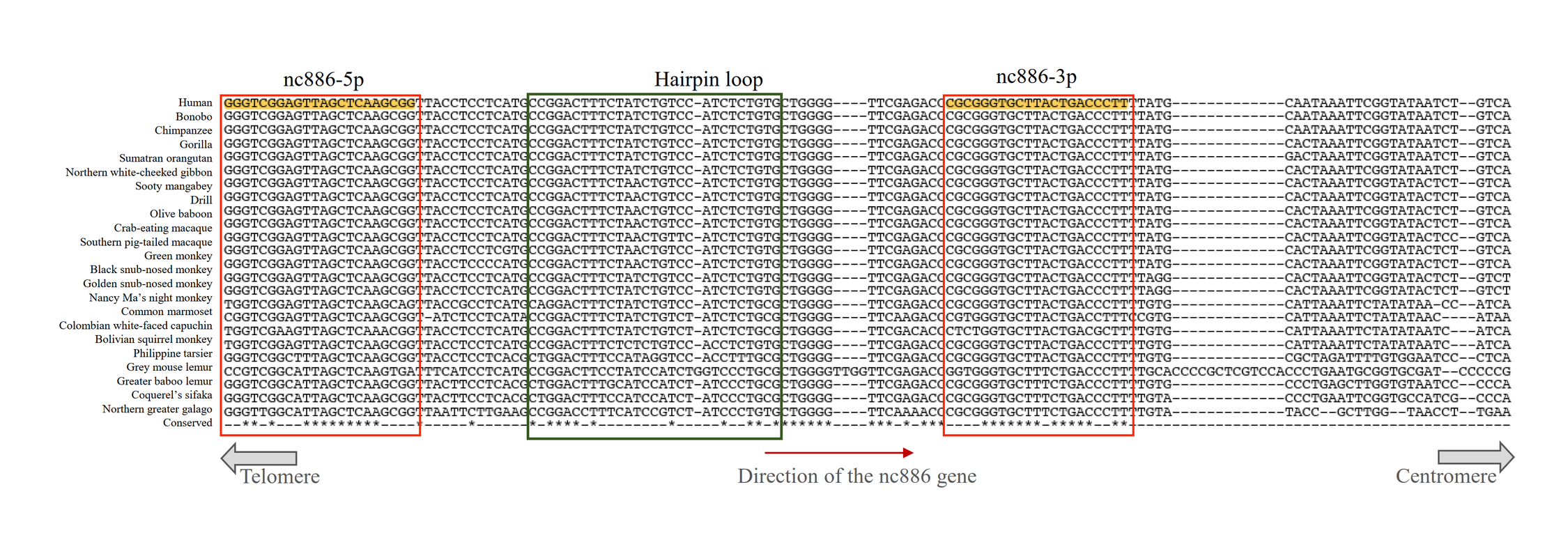

### S3 Figure

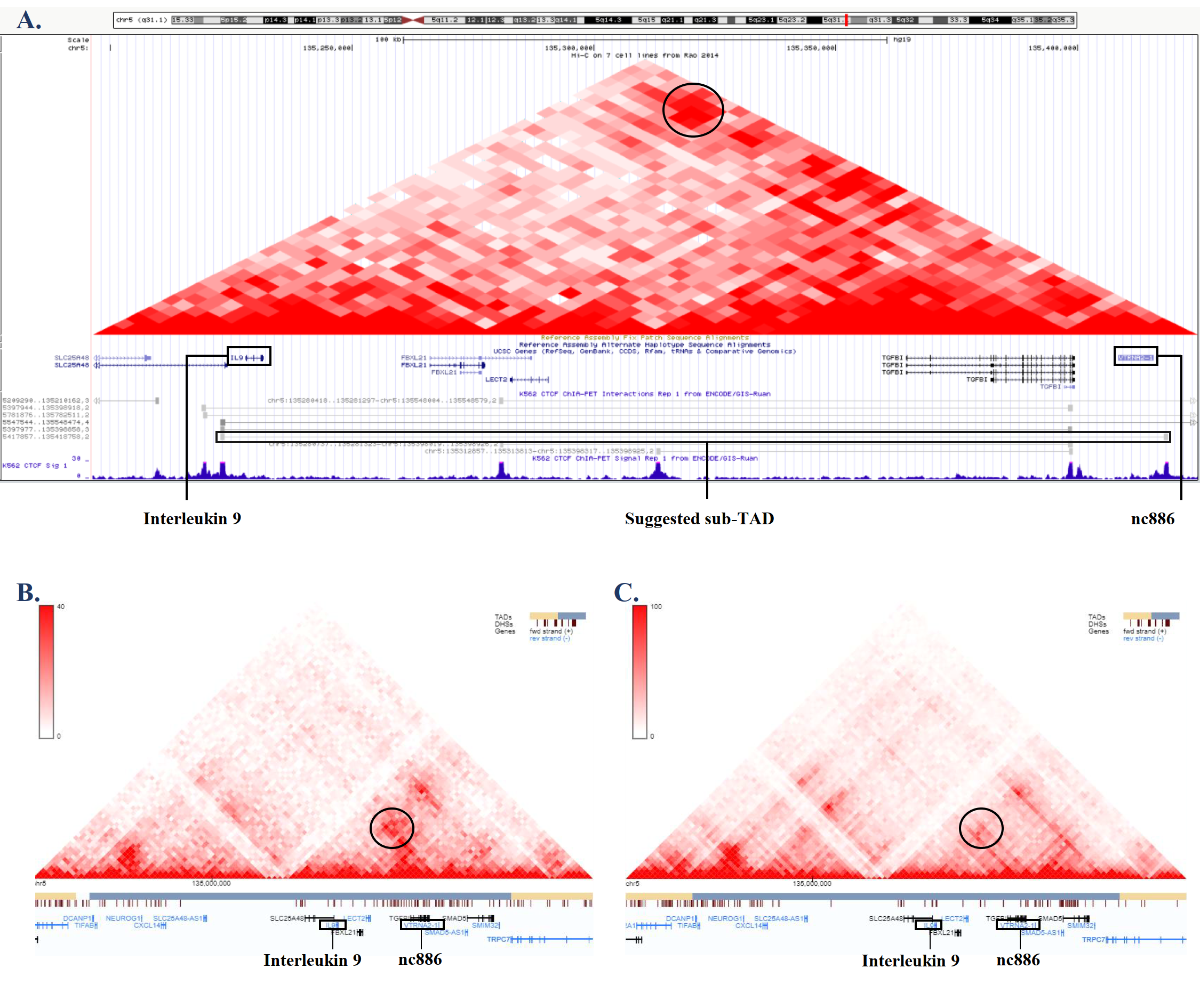

### S4 Figure

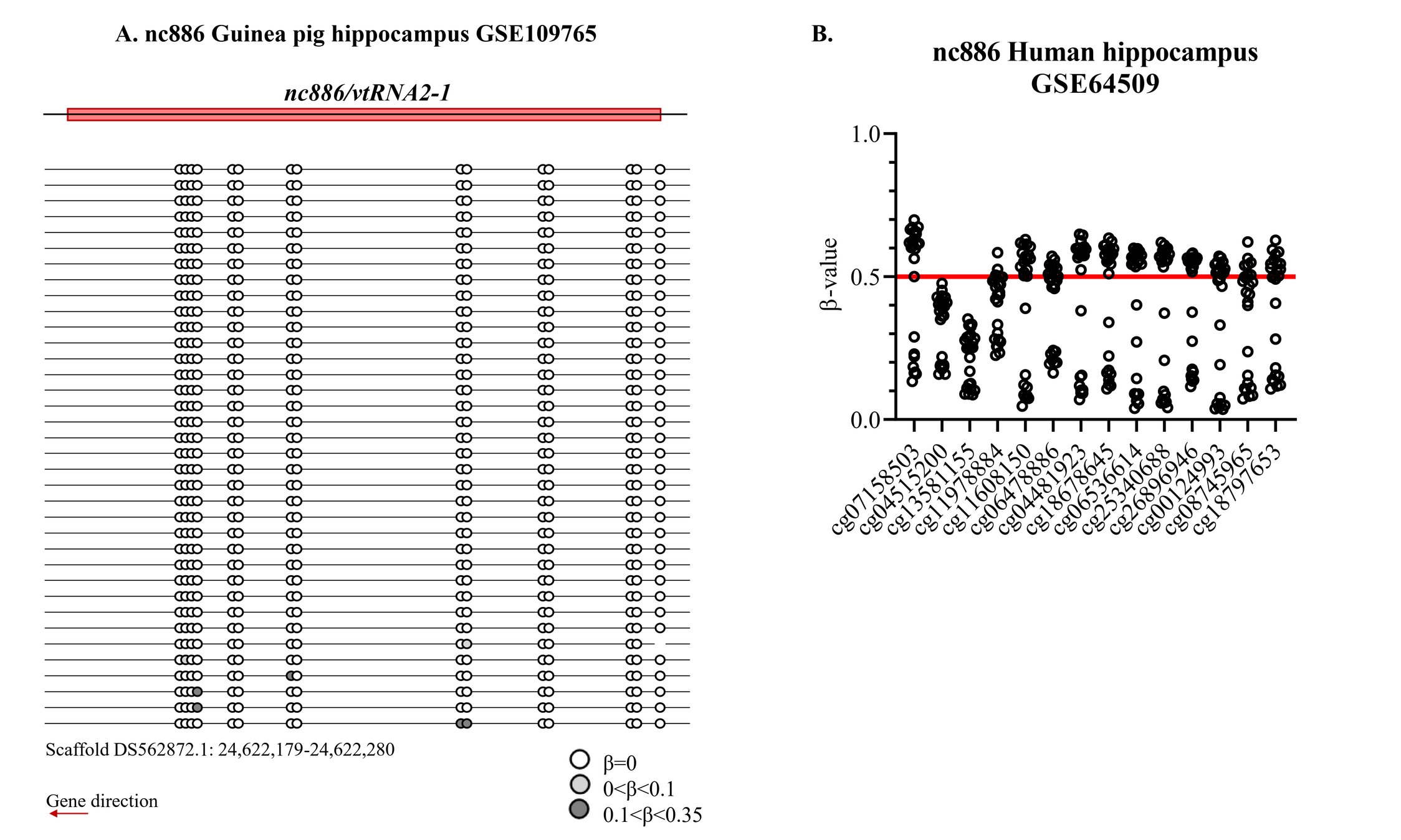

### S5 Figure

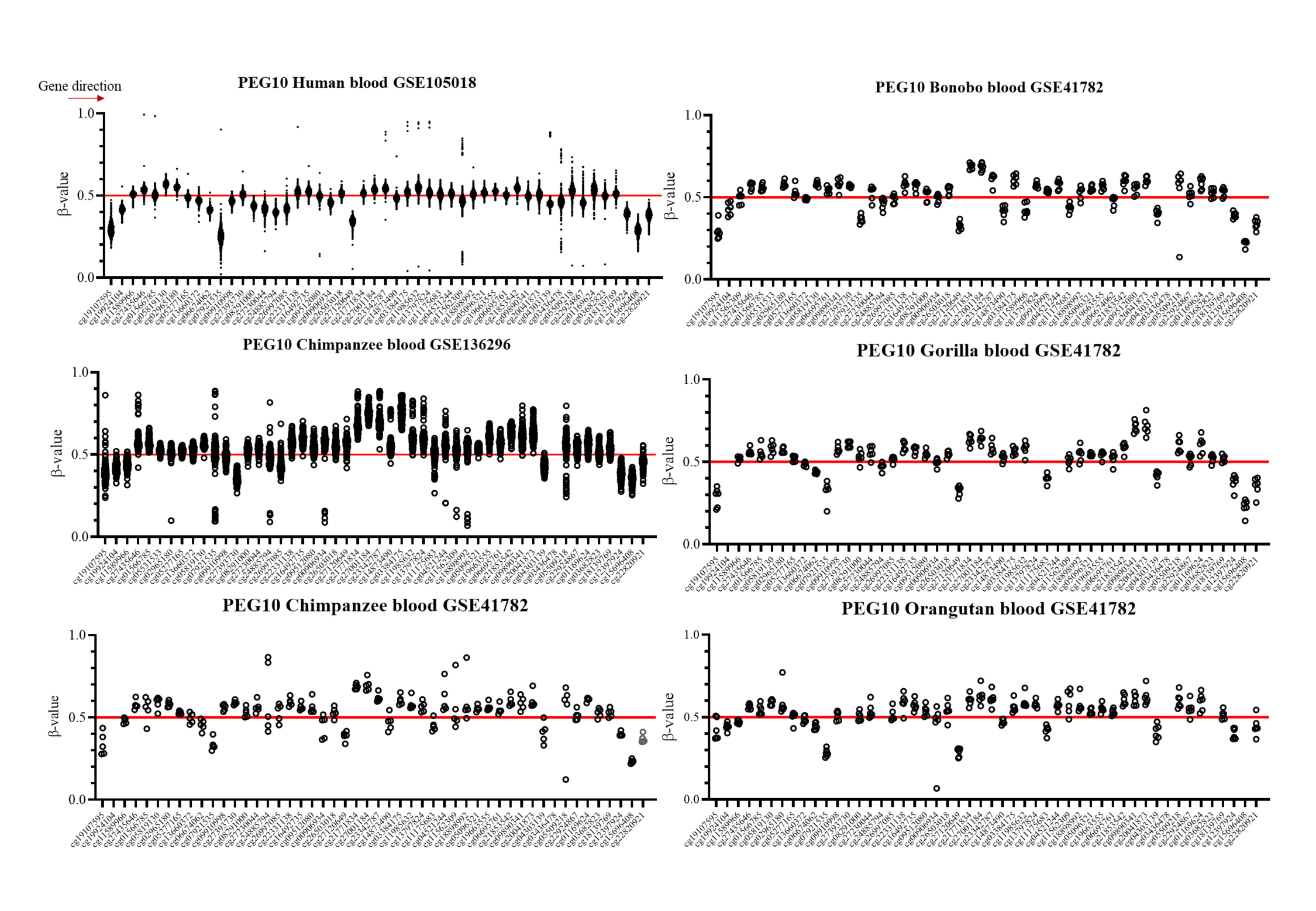

### S6 Figure

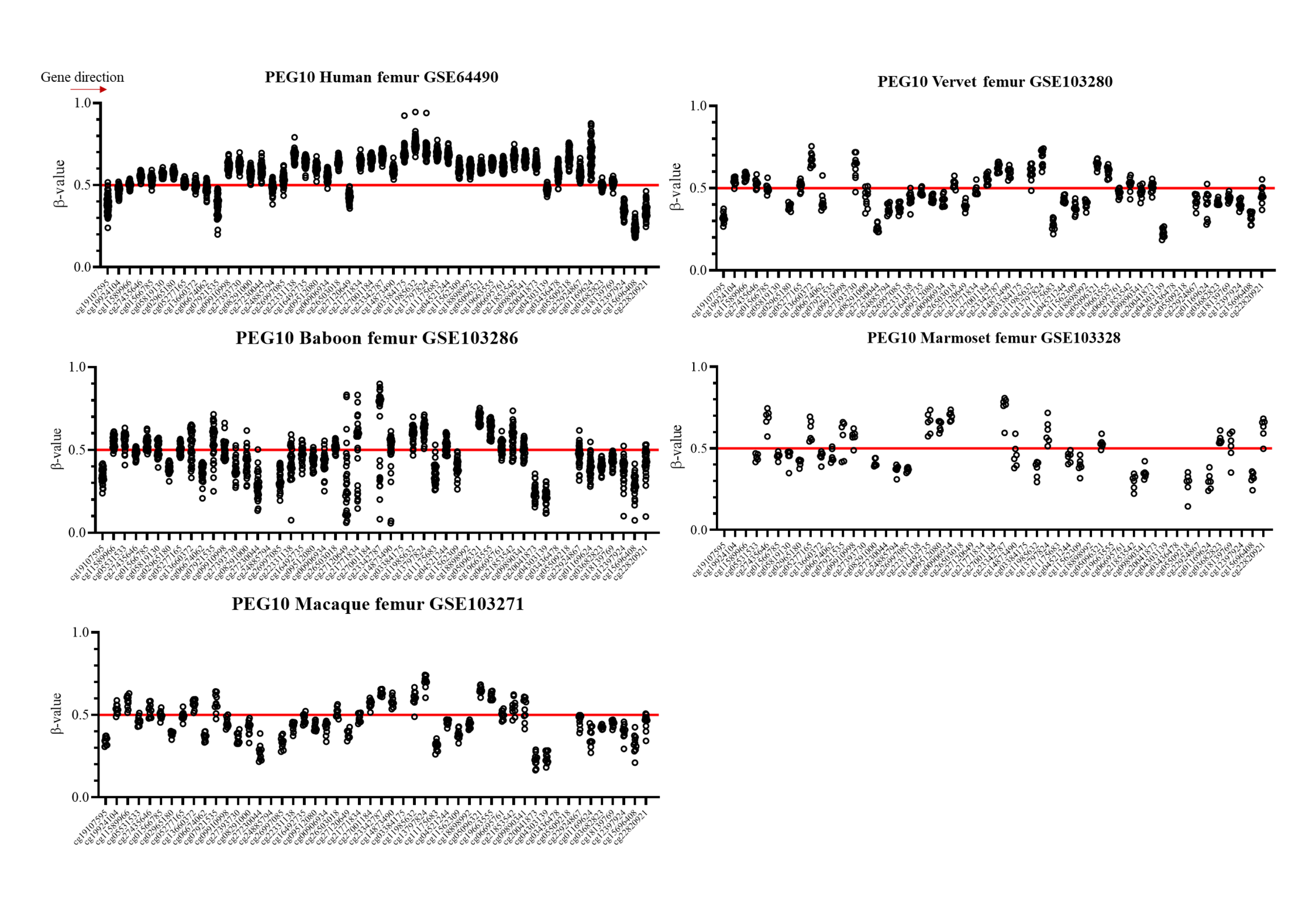
