## Supplementary material for "Methylation pattern of polymorphically imprinted *nc886* is not conserved across mammalia": S1 File

```
                                                                                                          cg07158503
Human       CTGTAAGTACAGCAAGGGGAGAGGCCCCTGCCCTTGTATGGCTTCTAGGGGGGAGCCCCCACGCTAAAGGTAAAGACAAATTGCATAAAATGTCTGGACACGATGCGAACAGGTGAGAAT
Bonobo      CTGTAAGTACAGCAAGGGCAGAGGCCCCTGCCCTTGTATGGCTTCTAGGGGGGAGCCCCCACGCTAAAGGTAAAGACAAATTGCATAAAATGTCTGGACACGATGCGAACAGGTGAGAAT
Chimpanzee  CTGTAAGTACAGCAAGGGCAGAGGCCCCTGCCCTTGTATGGCTTCTAGGGGGGAGCCCCCACGCTAAAGGTAAAGACAAATTGCATAAAATGTCTGGACACGATGCGAACAGGTGAGAAT
Gorilla     CTGTAAGTACAGCAAGGGCAGAGGCCTCTGCCCTTGTATGGCTTCTAGGGGGGAGCCCGCACGCTAAAGGTAAAGACAAATTGCATAAAATGTCTGGACACGATGCGAACAGGTGAGAAT
Orangutan   CTGTAGGTACAGCAAGGGCAGAGGCCCCTGCCCTTGTATGGCTTCTAGGGGGGAGCCCCCACGCTAAAGGTAAAGAAAAATTGCATAAAATGTCTGGACACGATGCGAACAGGTGAGAAC
Gibbon      CTGTAGGTACAACAAGGGCAGAGGCCCCTGCCCTTGTGTGGCTTCTAGGGGGGAGCCGCCACGCTAAAGGTAAAGAAAAATTGCATAAAATGTCTGGACACGATGCGAACAGGTGAGAAC
Baboon      CTGTGAGGACAGCAAGGGCAGACGCCTCTGCCCTTGTGCGGCTTCTAGCGGGGAGCCCCCA-ACTAAAGGAAAAGAAAAATTGCGTAAAATGTCTGGACACCATGCAAACAGGTGAGAAC
Macaque     CTGTGAGGACAGCAAGGGCAGACGCCTCTGCCCTTGTGCGGCTTCTAGCGGGGAGCCCCCACACTAAAGGAAAAGAAAAATTGTGTAAAATGTCTGGACACCATGCAAACAGGTGAGAAC
Vervet      CTGTGAGGACAGCAAGGGCAGAGGCCCCTGCCCTTGTGCGGCTTCCAGCGAAGAGTCCCCACACTAAAGGTAAAGAAAAATTGCATAAAAGGTGTGGACACGATGCAAACAGGTGAGAAC
Marmoset    CTGTGGGTACAGAAAAGGCAGACGCCTCTGCCCTTGTGCGGCTTCTAGAGGGGAAGCTCCACGCTAAAGGTAAAGAAAAATTGCATAAAATGTCTGGGCGCAATGCCAACAGATGAGAAC
            ****   * ***   ** ** *** *** ***********   ****** ** *   **   *   **    ******* ***** ******   ***** ** *** * * **** ***** ******

                                                                  cg04515200          cg13581155                          cg11978884
Human       AGCGTAGGTACAGGCCGCCAGGGAGGAAGAAACTTGGAACTTTCAAAGACGTAAAGAAAAATTCGGAGCGAGTCTCACTCGCTGTGGTCCAACAGGCAAGTCCTGGCGGGTGGCAATATA
Bonobo      AGCGTAGGTACAGGCCGCCAGGGAGGAAGAAACTTGGAACTTTCAAAGACGTAAAGAAAAATTCGGAGCGGGTCTCACTCGCTGTGGTCCAACAGGCAAGTCCTGGCGGGTGGCAATATA
Chimpanzee  AGCGTAGGTACAGGCCGCCAGGGAGGAAGAAACTTGGAACTTTCAAAGACGTAAAGAAAAATTCGGAGCGGGTCTCACTCGCTGTGGTCCAACAGGCAAGTCCTGGCGGGTGGCAATATA
Gorilla     AGCGTAGGTACAGGCCGCCAGGGAGGAAGAAACTTGGAACTTTCAAAGACGTAAAGAAAAATTCCTGGCGGGTCTCACTCGCTGTGGTCCAACAGGCAAGTCCTGGCGGGCGGCAATATA
Orangutan   AGCGTAGGTACAGGCCGCCAGGGAGGAAGAAACTTGGAACTTTCAAAGACGTAAAGAAAAATTCCGAGTGGGTCTCACTCGCTGTGGTCCAACAGGCAAGTCCTGGCGGGCGGCAATATA
Gibbon      AGCGTAGGCACAGGCCGCCAGCGAGGAAGAAACTTGGAGCTTTCAAAGACGTAAAGAAAAATTCCGAGCGGGTATCACTCGCTGTGGTCCACCAGGGAAGTCCTGGCGGGCGGCAATATA
Baboon      AGCGTAGGCACAGGCCACCAGGGAGGAAGAAACTTGGAGCTTTCAAAGATGTAAAGAAAAATTCCGAGCGGGTCTCACTTGCTGTGGTCCAACAGGCAAGTCCTGGTGGGCGGCAATATA
Macaque     AGCGTAGGCACAGGCCACCAGGGAGGAAGAAACTTGGAGCTTTCAAAGATGTAAAGAAAAATTCCGAGCGGGTCTCACTTGCTGTGGTCCAACAGGCAAGTCCTGGTGGGCGGCAATATA
Vervet      AGCGTAGGCACAGGCCACCAGGGAGGAAGAAACTTGGAGCTTTCAAAGATGTAAAGAAAAATTCCAAGCGGGTCTCACTCGCTGTGGTCCAACAGGCAAGTCCTGGTGGGCGGCAATATA
Marmoset    AGCACAGGCACAGGCCACCAGAGAGGAGGAAACTTGGAGCTTTCAAAAACATAAAGAAAAATTCCGAGGGGATCTCACTGGCTGTAGTCGAACAGGCAAGTCCTGGTGGGCGGCAATATA
            ***   *** ******* **** ***** ********** ******** *   *************    * *   * ***** ***** *** * **** ********* *** *********

                                                                                                                   cg11608150
Human       TGTGAAAAAAACAGTCATACAGATAAAACAGTAGACGGCAGCTATTGCTGTTAAACTTCCGTCCTACTTGGGACGAAATCCATAACGCACTCCGCGGTCCGAAGTTGTGGGGTCTCGGAG
Bonobo      TGTGAAAAAAACAGTCATACAGATAAAACAGTAGACGGCAGCTATTGCTGTTAAACTTCCGTTCTACTTGGGACGAAATCCATAACGCACTCCGCGGTCCAAAGTTGTGGGGTCTCGGAG
Chimpanzee  TGTGAAAAAAACAGTCATACAGATAAAACAGTAGACGGCAGCTATTGCTGTTAAACTTCCGTTCTACTTGGGACGAAATCCATAACGCACTCCGCGGTCCGAAGTTGTGGGGTCTCGGAG
Gorilla     TGTGAAAAAAACAGTCATACAGATAAAATAGTAGACGGCAGCTATTGCTGTTAAACTTCCGTTCTACTTGGGACGAAATCCATAACGCACTCCGCGGTCCGAAGTGGTGGGGTCTCGGAG
Orangutan   TGTAAAAAAAACAGTCATACAGATAAAATAGTAGACGGCAGCTATTGCTGTTAGACTTCCGTTCTACTTGGGACGAAATCCATAACGCACTCCGCGGTCCGAAGTTGTGGGGTCTCGGAG
Gibbon      TGTGAAGAAAACAGTCATACAGATAAAATAGTAGACTGCGGCTATTGCTGTTAAACTTCCGTTCTACTTGGGACGAAATCCATAACGCACTCCGCGGTCCGAAGTTGTGGGGTCTCGGAG
Baboon      TGTGAAGAAAACAGTCATACAGATAAAATAGTAGACGGCAGCTATTGCTGTTAAACTTCCACTCTACTTGGGACGAAATCTGTAACGCACTCCGCGGTCCGAGGTTGTGGGGTCTTGGAG
Macaque     TGTGAAGAAAACAGTCATACAGATAAAATAATAGACGGCAGCTATTGCTGTTAAACTTCCACTCTACTTGGGACGAAATCTGTAACGCATTCCGCGGTCCGAAGTTGTGGGGTCTTGGAG
Vervet      TGTGAAGAAAACAGTCGTACAGATAAAATAATAGACGGCAGCTATTGCTGTTAAACTTCCACTCTACTTGGGACGAAATCCGTAACGCACTCCGCGGTCCGAAGTTGTGGGGTCTTGGAG
Marmoset    TGCGAAATAAAAAGCTATACAGATAAAATAACAGACGGCAGCTATTGCCGTTAAACTTCCATTCTACTTGGAAGGAAATCTGTAATGCATTCCGTGACCCGAATTTGCGGGGTCTCAGAG
            **   **   *** **    *********** *   **** ** ******** **** ******   ******** * ******   *** *** **** *   ** *   * * *******   ***
```

```
                                                                                         cg06478886
Human      CTTCCAGGTGTGTCTCGTGAACTGCACAAGCATTTT-TTTGTCCCCATGCGTCCTACCTGGCAGTACAGGCTGGTCACGCACGCCCTGTAAGACCAGTGGCCAGCCTCCATACTTTCTGT
Bonobo     CTTCCAGGTGTGTCTCGTGAACTGCACAAGCATTTT-TTTGTCCCCATGCGTCCTACCTGGCAGTACAGGCCGGTCACGCACGTCCTGTAAGACCAGTGGCCAGCCTCCATACTTTCTGT
Chimpanzee CTTCCAGGTGTGTCTCGTGAACTGCACAAGCATTTT-TTTGTCCCCATGCGTCCTACCTGGCAGTACTGGCCGGTCACGCACGCCCTGTAAGACCAGTGGCCAGCCTCCATACTTTCTGT
Gorilla    CTTCCAGGTGTGTCTCGTGAACTGCACAAGCATTTT-TTTGTCCCCATGCGTCCTACCTGGCAGTACAGGCCGGTCACGCACGCCCTGTAAGACCAGTGGCCAGCCTCCATACTTTCTGT
Orangutan  CTTCCAGGTGTGTCTCGTGAACTGCACGAGCGTTTT-TTTGTCCCCATGCGTCCTACCTGGCAGTACAGGCCGGTCACGCACGCCCTGTAAGACCAGTGGCCAGCCTCCATACTTTCTGT
Gibbon     CTTCCAGGTGTGTCTCGTGAACTGCACGAGCATTTT-TTTGTCCCCATGCATCCTACCTGGCAGTAGAGGCCGGTCACGCAAGCCCTGTAAGACCAGTGGCCCACCTCCATACTTTCTGT
Baboon     CTTCAAGGTGTGTCTCGTAAACTGCACGAGCATTTT-TTTCTTCCCCTGCATCCCACCTGGCAGTACAGGCCGGTCACGCACGCCCTGTAAGACCAGTGGCCCACCTCCATACTTTCTGT
Macaque    CTTCAAGGTGTGTCTCGTGAACTGCACGAGCATTTTTTTCTCCCCCTGCGTCCCACCTGGCAGTACAGGCCGGTCACGCACTCCATGTAAGACCAGTGGCCCACCTCCATACTTTCTGT
Vervet     CTTCAAGGTGTGTCTCGTGAACTGCACGAGCATTTT-TTTCTGCCCCTGCATCCCACCTGGCAGTACAGGCCGGTCACGCACGCCCTGTAAGACCAGTGGCCCATCTCCATACTTTCTGT
Marmoset   TTTCAAGGTGTGTCTCGTGAACTGCACGAGCATTTT-TTTGTCCCCATGCATCCTACCTGGCAGTAGAGGCCGGTCACACCCCCCCCATAAGACCAGTGGGCCACCTCCATAATTTCTGT
            *** ************* ******** *** **** *** * *** *** *** ************ *** ****** *        ************     ******* *******

Human      CACACCTTCAAAGTGACACCAACTTATGTTATCAGCCTATTAATTTAGCAAAGTTAAAAGGGATAAAAAATA---ACAAAGCCTTCAGAGTGACAGATTATACCGAATTTATTGCATAAA
Bonobo     CACACCATCAAAGTGACACCAACTTATGTTATCAGCCTATTAATTTAGCAAAGTTAAAAGGGATAAAAAATA------AAGCCTTCAGAGTGACAGATTATACCGAATTTATTGCATAAA
Chimpanzee CACACCATCAAAGTGACACCAACTTATGTTATCAGCCTATTAATTTAGCAAAGTTAAAAGGGATAAAAAATAACA---AAGCCTTCAGAGTGACAGATTATACCGAATTTATTGCATAAA
Gorilla    TACACCATCAAAGTGACACCAACTTATGTTATCAACCTATTAATTTAGCAAAGTTAAAAGGGATAAAAAATA---ACAAAGCCTTCAGAGTGACAGATTATACCGAATTTAGTGCATAAA
Orangutan  CACACCATCAAAGCGACACCAACTTATGTTATCAACCTATTAATTTAGCAAAGTTAAAAGGGACAAAAAATA---ACAAAGCCTTCAGAGTGACAGATTATACCGAATTTAGTCCATAAA
Gibbon     CATACCATCAAAGCGACACCAACTTATGTTATCAACCTATTAATTTAGCAAAGTTAAAAGGGACGAAAGATA---ACAAAGCCTTCAGAGTGACAGATTATACCGAATTTAGTGCATAAA
Baboon     CGCACCATCAAAGCGACACCAACTTATGTTATCAACCTATTAATTTAGCAAAGTTAAAAGAGACAAAAAGTA---ACAAAGTCTTCAAAGTGACAGAGTATACCGAATTTAGTGCATAAA
Macaque    CGCACCATCAAAGCGACACCAACTTATGTTATCAACCTATTAATTTAGCAAAGTTAAAAGAGACAAAAAGTA---ACAAAGTCTTCAAAGTGACGGAGTATACCGAATTTAGTGCATAAA
Vervet     CGCACCATCAAAGCGGCACCAACTTATGTTATCAACCTATTAATTTAGCAAAGTTAAAAGGTATACCGAATTTAGTGCATAAAAGGGTCAGTAAGCACCCGCGGGTCAGACAAAAAATA-
Marmoset   TGCACTATTAAAGCAACACCAATTTACATTATCCGCCTATTAATGTAGCAAAATTAAAAGTTATATAGAATTTAATGCACGGAAAGGTCAGTAAGCACCCACGGGCCGAAAAAAAACC-
              ** * ****   ****** ***  *****  ********* ******* *******                                *   *

                cg04481923
Human      AGGGTCAGTAAGCACCCGCGGGTCTCGAACCCCAGCACAGAGATGGACAGATAGAAAGTCCGGCATGAGGAGGTAACCGCTTGAGCTAACTCCGACCCGGGTAGGAGTGTGCGACTGAAA
Bonobo     AGGGTCAGTAAGCACCCGCGGGTCTCGAACCCCAGCACAGAGATGGACAGATAGAAAGTCCGGCATGAGGAGGTAACCGCTTGAGCTAACTCCGACCCGGGTAGGAGTGTGCGACTGAAA
Chimpanzee AGGGTCAGTAAGCACCCGCGGGTCTCGAACCCCAGCACAGAGATGGACAGATAGAAAGTCCGGCATGAGGAGGTAACCGCTTGAGCTAACTCCGACCCGGGTAGGAGTGTGCGACTGAAA
Gorilla    AGGGTCAGTAAGCACCCGCGGGTCTCGAACCCCAGCACAGAGATGGACAGATAGAAAGTCCGGCATGAGGAGGTAACCGCTTGAGCTAACTCCGACCCGGGTAGGAGTGTGCGACTGAAA
Orangutan  AGGGTCAGTAAGCACCCGCGGGTCTCGAACCCCAGCACAGAGATGGACAGATAGAAAGTCCGGCATGAGGAGGTAACCGCTTGAGCTAACTCCGACCCGGGTAGGAGTGTGCGACTGAAA
Gibbon     AGGGTCAGTAAGCACCCGCGGGTCTCGAACCCCAGCACAGAGATGGACAGATAGAAAGTCCGGCATGAGGAGGTAACCGCTTGAGCTAACTCCGACCCGGGTAGGAGTGTGCGACTGAAA
Baboon     AGGGTCAGTAAGCACCCGCGGGTCTCGAACCCCAGCACGGAGATGGACAGTTAGAAAGTCCGGCATGAGGAGGTAACCGCTTGAGCTAACTCCGACCCGGTTAGAAGTGTGCGATTGAAA
Macaque    AGGGTCAGTAAGCACCCGCGGGTCTCGAACCCCAGCACAGAGATGAACAGTTAGAAAGTCCGGCATGAGGAGGTAACCGCTTGAGCTAACTCCGACCCGGTTAGAAGTGTGCGATTGAAA
Vervet     --ACAAAGTCTTCAAAGTGACAGATCGAACCCCAGCACAGAGATGGACAGTTAGAAAGTCCGGCATGGGGAGGTAACCGCTTGAGCTAACTCCGACCCGGTTAGAAGTGTGCGATTGAAA
Marmoset   --ATAAAGTCCTCAAAGTTATG--TTGAACCCCAGCGCAGAGATAGACAGATAGAAAGTCCGGTATGAGGAGATA-CCGCTTGAGCTAACTCCGACCGGTATAGGGGTGTGCAGTTGAAA
                  **   **          * ********** *******  **** ************ *** **** ** ********************* *  ***  ******   *****
```

```
                                  cg26896946
           cg18678645                                                  cg06536614        cg25340688   |   cg00124993
Human      CTTCTAAACCATAGAAGAGTGACGATGTGGAGAGGGACGGGCTGCATGTGCTCCCCGCCCCCGAGAGGCCTGCGTCATGCGGTCTCGCCCGCTCTGCGCCAGGCGTCCTGCTAACGTGTC
Bonobo     CTTCTAAACCATAGAAGAGTGACGATGTGGAGAGGGACGGGCTGCATGTGCTCCCCGCCCCCGAGAGGCCTGCGTCATGCGGTCTCGCCCGCTCTGCGCCAGGCGTCCTGCTAACGTGTC
Chimpanzee CTTCTAAACCATAGAAGAGTGACGATGTGGAGAGGGACGGGCTGCATGTGCTCCCCGCCCCCGAGAGGCCTGCGTCATGCGGTCTCGCCCGCTCTGCGCCAGGCGTCCTGTTAACGTGTC
Gorilla    CTTCTAAACCATAGAAGAGTGACGACGTGGAGAGGGACGGGCTGCATGTGCTCCCCGCCCCCGAGAGGCCTGCGTCATGCGGTCTCGCCCGCTCTGCGCCAGGCGTCCTGCTAACGTGTC
Orangutan  CTTCTAAACCATAGAAGAGTGACGACGTGGAGAGGGACGGGCTGCATGTGTTCCCCGCCCCCGAGAGGCCTGCGTCATGCGGTCTCGTCCGCTCTGCGCCAGGCGTCCTGCTAACGTGTC
Gibbon     CTTCTAAACCATGGAAGAATGACGACGTGGAGAGGGACGGGCTGCATGTGTTCCCCGCCCCCGAGAGGCCTGCGTCATGCGGTCCCGCTCGCTCTGCACCAGGCGTCCTGCTAACGTGTC
Baboon     CTTCTAAACCATAGAAGAGTGACGACGTGGAGAGGGACGGGCTGCATGTGCTCCCCGCCCCCAAGAGGCCTGCGTCATGCGGTCCCGCCCGCTCTGCGCCAGGCGTCCTGCTAGCGTGTC
Macaque    CTTCTAAACCATAGAAGAGTGACGACGTGGAGAGGGACGGGCTGCATGTGCTCCCCGCCCCCAAGAGGCCTGCGTCATGCGGTCCCGCCCGCTCTGCGCCAGGCGTCCTGCTAGCGTGTC
Vervet     CTTCTAAACCATAGAAGAGTGACGACGTGGAGAGGGACGGGCTGCATGTGTTCCCCGCCCCCAAGAGGCCTGCGTCATGCGGTCCCGCCCGTTCTGCGCCAGGCGTCCTGCTAGCGTGTC
Marmoset   CTTCTAAACCATAGAAGAGTGACAGCGTGGAGAGGGAAGGGCTGCATGTGTTCCCCACCCCCGAGAGGCCTGCGTCACGTGGTCCCGCCAGCTCTGCGCCAGGCGTCCTGCTAGCGTGAC
           ****************** ****   *********** ************ ***** ***** ************** * **** **    * ***** ************ ** **** *

                                                                                                                  cg08745965
Human      CTGGAGGGACTCTCAGTTCCTCCCGCCC-GCATCCTGCGCGGGAACCGTGGAAGGGGGCAAAATCCACCCACTGGAGGGGAGGC-AGGAGGGTGCGGGGGGGCGTGTGGGCCGTCTACCT
Bonobo     CTGGAGGGACTCTCAGTTCCTCCCGCCC-GCATCCTGCGCGGGAACCGTGGAAGGGGGCAAAATCCACCCACTGGAGGGGAGGC-AGGAGGGTGTGGGGGGGCGTGTGGGCCGTCTACCT
Chimpanzee CTGGAGGGACTCTCAGTTCCTCCCGCCC-GCATCCTGCGCGGGAACCGTGGAAGGGGGCAAAATCCACCCACTGGAGGCGAGGC-AGGAGGGTGCGGGGGGGCGTGTGGGCCGTCTACCT
Gorilla    CTGGAGGGACTCTCAGTTCCTCCCGCCC-GCATCCTGCGCGGGAACCGTGGAAGGGGGCAAAATCCACCCACTGGAGGGGAGGC-AGGAGGGTGCGGGGGGGCATGTGGGCCGCCTACCT
Orangutan  CTGGAGGGCCTCTCAGTTCCTCCCGCCC-GCATCCTGCGCGGGAACCGTGGAAGGGGGCAAAATCCACCCACTGGAGGGGAGGC-AGGAGGGCGCAGGGGGGCATGTGGGCCGCCTACCT
Gibbon     CTGGAGGGCCTCTCAGTTCCTCCCGCCC-GCATCCTGCGCGGGAACCGTGGAAGGGGGCAAAATCCACCCACTGGAAGGGAGGC-AGGAGGGCGCAGGG-GCCATGTGGGCCGCCTACCT
Baboon     CTGGAGGGCCTCTCGGTTCCTCCCGCCC-CCATCCTGCGCAGGAACCGTGGAAGGGGGCACAATCCACCCA--GGAGGGGAGGC-ACGAGGGCGCGGGGGAACATGTGGGCCGCCTACCT
Macaque    CTGGAGGGCCTCTCGGTTCCTCCCGCTCCCCATCCTGCGCAGGAACCGTGGAAGGGGGCACAATCCAC-CGCAGGAGGGGAGGCGACGAGGGCGCGGGGGAACATGTGGGCCGCCTACCT
Vervet     CTGGAAGGCCTCTCAGTTCCTCCCGCTC-CCATCCTGCGCAGGAACCGTGGAAGGGGGCACAATCCAT-CGCAGGAGGGGAGGCGACGAGGGCGCAGGGGAACATGTGGGCCGCCTACCT
Marmoset   CTGGAGATCCTCCCAGTTCCTCCCGCCC-CGATCCAGCGCAGGAACTGTGGAAGAGGGCAAAATACAATC-CTGGAGGGGAAGC-AGGAGGGCACGGGGAAGCATGCGGGCTGCCTACCT
           *****    *** ************* *   **** **** ***** ******* ***** *** **  *    *** * ** ** * *****    ***   * ** **** * ******

                                                                                cg18797653
Human      AGGTCCAGCAGCCAGGCTGCTGAGGAGTACCCCCGCCAAAGGCTTTTCGGGGTTCTCTCCAGCAGACGGGGGCAGCCTAAGGCCTCCATAAAATCTCCCCGAAGCAGCCTATGAACTTGC
Bonobo     AGGTCCAGCAGCCAGGCTGCTGAGGAGTACCCCCGCCAAAGGCTTTTCGGGGTTCTCTCCAGCAGACGGGGGCAGCCTAAGGCCTCCATAAAATCTCCCCGAAGCAGCCTATGAACTTGC
Chimpanzee AGGTCCAGCAGCCAGGCTGCTGAGGAGTACCCCCGCCAAAGGCTTTTCGGGGTTCTCTCCAGCAGACGGGAGCAGCCTAAGGCCTCCATAAAATCTCCCCGAAGCAGCCTATGAACTTGC
Gorilla    AGGTCCAGCAGCCAGGCTGCTGAGGAGTACCCCCGCCAAAGGCTTTTCGGGGTTCTCTCCAGCAGACGGGGGCAGCCTAAGGCCTCCATAAAATCTCCCCGAAGCAGCCTATGAACTTGC
Orangutan  AGGTCCAGCAGCCAGGCTGCTGAGAAGTACCCCCGCCAAAGGCTTTTCGGGGTTCTCTCCAGCAGACGGGGGCAGCCTAAGGCCTCCATAAAATCTCCCCGAAGCAGCCTATGAACTTGC
Gibbon     AGGTCCAGCAGCCAGGCTGCCGAGGAGTACCCCCGCCAAAGGCTTTTCGGGGTTCTCTCCAGCAGACGGGGGCAGCCTAAGGCCTCCATAAAATCTCCCCGAAGCAGCCTATGAACTTGC
Baboon     AGGTCCAGCAGCCAGGCTGCTGAGGAGTACCGCCGCCAAGGGCTTTTTGGGGTTCTCTCCAGCAGACCGGGGCAGCCTAAGGCCTCCATAAAA-CTCCCCCAAGCAGCCTATGAACTTGC
Macaque    AGGTTCAGCAGCCAGGCTGCTGAGGAGTACCGCCGCCAAGGGCTTTTTGGGGTTCTCTCCAGCAGACCGGGGCAGCCTAAGGCCTCCATAAAACTGCCCCCAAGCAGCCTATGAACTTGC
Vervet     AGGTCCAGCAGCCAGGCTGCTGAGGAGTACCACCGCCAAAGGCTTTTTGGGGTTCTCTCCAGCAGACCGGGGCAGCCTAAGGCCTCCATAAAACTGCCCCCAAGCAGCCTATGAACTTGC
Marmoset   AGGTCTGGCAGCCAGGCTGCTGAGGAGTACCCCCGC-AAAGGCTTTTTGGGGTTCCCTCCAGCAGACCAGGGCAGTCTAAGTCCTCCATAAAATCACCCCCAAGCAGCCTGTGAACTTGC
           ****   ************* *** ****** **** ** ******* ******* ***********  * **** ***** ***********   **** ********* *********
```
